# Do-it-yourself *de novo* antibody sequencing workflow that achieves complete accuracy of the variable regions

**DOI:** 10.1101/2024.12.23.630060

**Authors:** Meng-Ting He, Ning Li, Jian-Hua Wang, Zhi-Zhong Wei, Jie Feng, Wen-Ting Li, Jian-Hua Sui, Niu Huang, Meng-Qiu Dong

## Abstract

Antibodies are widely used as research tools or therapeutic agents. Knowing the sequences of the variable regions of an antibody—both the heavy chain and the light chain—is a prerequisite for the production of recombinant antibodies. Mass spectrometry-based *de novo* sequencing is a frequently used, and sometimes the only approach to gaining this information. Here, we describe a workflow that enables accurate sequence determination of monoclonal antibodies based on mass spectrometry data and freely available software tools. This workflow, which we developed using a homemade anti-FLAG monoclonal antibody as a reference sample, achieved 100% accuracy of the variable regions with clear distinction between leucine (L) and isoleucine (I). Using this workflow, we successfully decoded a monoclonal anti-HA antibody, for which we had no prior knowledge of its sequence. Based on the *de novo* sequencing result, we generated a recombinant anti-HA antibody, and demonstrated that it has the same specificity, sensitivity, and affinity as the commercial antibody.

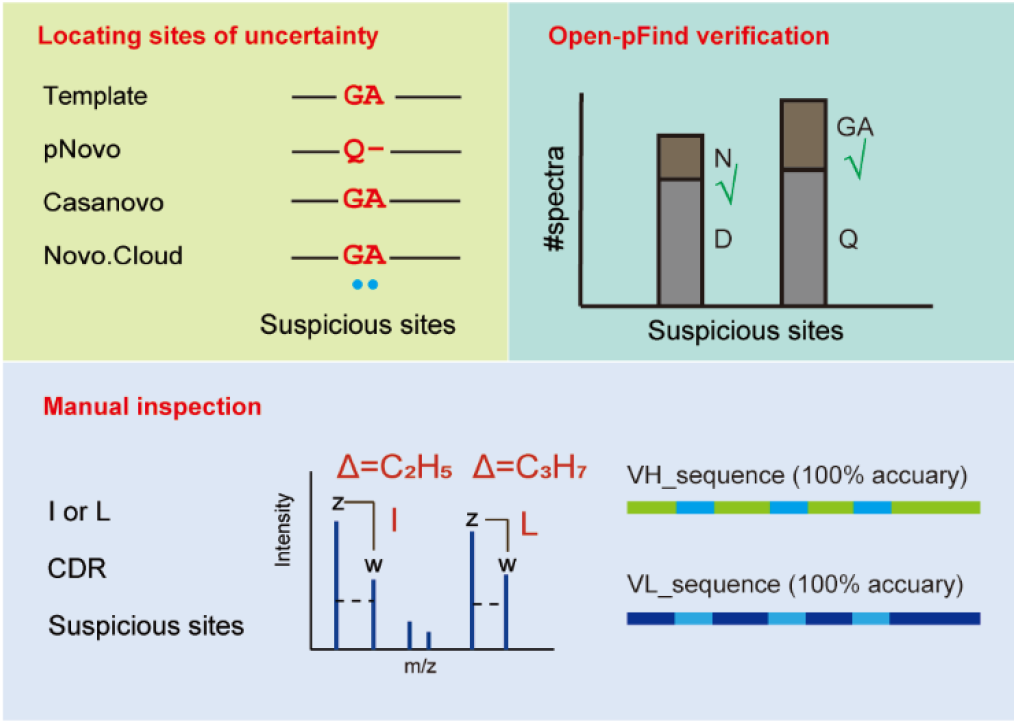

## INTRODUCTION

Antibodies are important both as therapeutic agents in medicine and as research tools in life science^1, 2^. There is a growing preference for recombinant antibodies in long-term studies due to their batch-to-batch reproducibility, animal-free manufacturing, and economic advantages^3, 4^. The sequences of the variable light (VL) and variable heavy (VH) chains must be known in order to produce or engineer recombinant antibodies. For instance, humanized monoclonal antibodies, achieved by grafting complementarity-determining region (CDR), have clear advantages in safety and effectiveness as therapeutic antibodies^5-7^. The ability to manipulate antibody sequences creates new opportunities for enhancing their clinical utility.

Hybridoma sequencing, i.e., sequencing of RNA from hybridoma cells, is the primary method for antibody sequencing^8, 9^. However, this method is not always feasible, for example, when the cell line is lost or when the antibody comes from an unreplenishable source, for example, patient serum. Additionally, hybridoma cells may mutate, causing deviations in VL or VH sequences if left unchecked. In these cases, *de novo* antibody sequencing offers an alternative. This approach does not rely on hybridoma cells or genetic materials. It involves digesting a purified antibody into peptides, analyzing these peptides via liquid chromatography coupled with tandem mass spectrometry (LC-MS/MS), identifying their sequences from the resulting MS data, and finally assembling the peptide sequences to reconstruct the full antibody sequence.

Advancement of software tools have made *de novo* antibody sequencing a reality. Among these are various *de novo* sequencing software tools, such as PEAKS^10^, PepNovo+^11^, pNovo^12^, Novor^13^, DeepNovo^14^, SMSNet^15^, MaxNovo^16^, PGPointNovo^17^, Casanovo^18^, BiATNNovo^19^, and π-HelixNovo^20^. Also, there are tools for antibody sequence assembly such as Meta-SPS^21^, pTA^22^, MuCS^23^, and Stitch^24, 25^. Commercial software packages, including Rapid Novor, PEAKs AB^26^, and Supernovo^27^ have gained popularity for their user-friendly interfaces and robust assembly of peptide fragments, thanks in part to the use of antibody germline sequences as reference.

Despite the remarkable progress, *de novo* antibody sequencing still faces challenges:

1. *de novo* peptide sequencing results are often inaccurate. Bealie et al^28^. conducted a comprehensive evaluation of several *de novo* peptide sequencing tools, including Novor, pNovo3, DeepNovo, SMSNet, PointNovo, and Casanovo, alongside the assembler de Bruijn ALPS. The presence of various types of error hinders severely the effectiveness of the assembly software. Sequence scrambling within the first or last three amino acids of a peptide (e.g. PGS to SPG, GPS or PSG) is particularly troublesome because it obstructs identification of overlapping sequences between peptides, which are critical for building a contig. Another common error concerns the ambiguity between N and GG, or among Q, AG, and GA, because they are of the same mass^25^. If fragment ions resulting from the cleavage between GG, AG, or GA are absent from some or all of the MS2 spectra of a peptide in question, then *de novo* sequencing software may place N at the position of GG or AG/GA.
2. Difficulty in isomeric amino acids I and L.
3. Challenges in *de novo* sequencing of peptides with post-translational modifications (PTMs). *De novo* sequencing algorithms typically operate with a predefined set of PTMs. Therefore, if a peptide carries a PTM outside the predefined PTM library, it will not be identified or could be misidentified as some other peptide. This will reduce the overall effectiveness of MS-based antibody sequencing. Worth noting is deamidation, a common PTM. Deamidated asparagine (N) is indistinguishable from aspartic acid (D), and similarly, deamidated glutamine (Q) is indistinguishable from glutamic acid (E). As a result, the presence of deamidation often leads to misassignment of D or E to where N or Q is, respectively.

In this study, we chose the software Stitch to assemble the relatively short *de novo* sequence reads into full-length antibody sequences. Stitch is available to the public free of charge and it has demonstrated success in sequencing an isolated Fab from patient serum, as well as in profiling the IgG repertoire of patients^24^. Peptide Tag Assembler (pTA)^22^ and Multiple Contigs & Scaffolding (MuCS)^23^ are two other free software tools built for similar purposes. Whereas pTA and MuCS rely almost entirely on overlapping *de novo* reads to build longer, contiguous sequences, Stitch additionally makes use of the antibody sequences deposited in the international ImMunoGeneTics (IMGT) information system or any user-defined antibody sequences as templates to place *de novo* peptide reads in the correct framework of the heavy chain and the light chain. Stitch also consults with the principle behind the genesis of an antibody sequence, i.e. somatic recombination of the Variable, Diversity (for the heavy chain), Joining, and Constant segments. Following the principle of V-(D)-J-C recombination, Stitch recombines the top-scoring V/J/C segments to produce new templates for a second round of template matching with the *de novo* reads. The sequence of CDR3, which lies in the V-(D)-J junction, is reconstructed by Stitch with overlapping *de novo* reads that extend out of the V- and J-segments. As such, Stitch is an effective antibody sequence assembler and is less susceptible to errors in *de novo* sequence reads than pTA and MuCS.

In this research, we introduce an effective and economical workflow for sequencing monoclonal antibodies. We applied this workflow to decode the sequence of an unknown anti-HA antibody and generated a recombinant antibody based on the antibody sequence thus obtained. The specificity, sensitivity, and affinity of this recombinant anti-HA antibody are indistinguishable from the original commercial antibody.

## EXPERIMENTAL SECTION

### Materials and reagents

Homemade anti-FLAG antibody was provided by Biologics Research Center & Antibody Center, NIBS, Beijing, China. Anti-HA antibody (TANA2) was purchased from MBL (# M180-3). PNGase F was purchased from NEB (#0704S). Trypsin (#v5280), Elastase (#v1891), Pepsin(#v1959), Chymotrypsin (#v1062) and Asp-N (#v1621) were purchased from Promega. GluC (#90054) was purchased from Thermo Fisher Scientific.

### Sample Preparation

5 μg sample was deglycosylated with PNGase F and then denatured in SDS loading buffer (95 °C, 10 min). Heavy chain and light chain were separated by gel electrophoresis and excised separately from the gel into 1 mm^3 pieces. The pieces after reducted by 10 mM DTT and alkylated by 55 mM IAA, finally individually digested with the following enzymes: Trypsin, Elastase, Pepsin, Chymotrypsin, Asp-N, GluC, Trypsin + AspN, Trypsin + GluC^29^.

### LC-MS/MS

The digested peptides were analyzed using an EASY-nLC 1200 system (Thermo Fisher Scientific) interfaced with a Fusion Lumos mass spectrometer (Thermo Fisher Scientific). Peptides were loaded onto a pre-column (75 μm ID, 5 cm long, packed with ODS-AQ 12 nm–10 μm beads) and separated on an analytical column (75 μm ID, 15 cm long, packed with Luna C18 1.9 μm 100 Å resin). Samples were separated with a 90 min linear reverse-phase gradient at a flow rate of 400 nL/min as follows: 6%-37% B in 79 min, 37%-100% B in 2 min, 100% B for 9 min (A = 0.1% FA, B = 80% ACN, 0.1% FA). Spectra were acquired in data-dependent mode: the top 15 most intense precursor ions from each full scan (resolution 60,000) were isolated for HCD or EThcD MS2 (resolution 30,000, HCD Collision Energy 30%) with a dynamic exclusion time of 30 s. Precursors with 1+, more than 7+, or unassigned charge states were excluded.

### MS Data Analysis

Firstly, raw data were converted to MGF files by pParse2.0, and then searched by pNovo (v.3.1.4, http://pfind.org/software/pNovo/), Novor.Cloud (https://app.novor.cloud/), Casanovo (v.3.5.0, https://github.com/Noble-Lab/casanovo) while raw data were used as input for Maxnovo (v.2.4.10.0, https://www.maxquant.org/). The parameters are set as follows: precursor tolerance was 10 ppm, fragment tolerance was 20 ppm. Carbamidomethylation (+57.021 Da) on cysteine was set as a static modification; oxidation of methionine (+15.995 Da); deamidation of asparagine (+0.984 Da), deamidation of glutamine (+0.984 Da) were set as variable modifications. The Casanovo_massivekb.ckpt model was utilized for mass spectrometry data of tryptic peptides, while the Casanovo_non-enzy.ckpt model was applied to mass spectrometry data of peptides digested by other enzymes. Secondly, the results of *de novo* peptide sequencing as input for assembly software Stitch (v1.5.0, https://github.com/snijderlab/stitch/releases/tag/v1.5.0), setting the cutoff score separately as 50, 60, 70, 80, 90 of pNovo, Novor.Cloud, MaxNovo, and 0, 0.2, 0.4, 0.6, 0.8, 0.9 of Casanovo.

### Sequence Calibration

Common confusion between same mass amino acids GA/Q, GG/N, N [Deamidation]/D, and Q[Deamidation]/E were compared with the template sequence. Amino acids that differ from the template sequence were marked as suspicious sites. The comparison of the antibody sequences obtained with different *de novo* sequence software by Muscle (https://github.com/rcedgar/muscle/releases/tag/5.1.0) and SnapGene could display the results, which differed from each other were also labeled as suspicious sites. sequence calibrations were analyzed by pFind3 (v3.2.0, http://pfind.org/software/pFind/) software in open search mode. MS Data Format was set to MGF, the consensus and template sequences from the output of Stitch were used as the database. The enzyme was selected as NoEnzyme, Non-specific. Precursor Tolerance was set to ±10 ppm, Fragment Tolerance was set to 20 ppm^30^. The number of support spectra of different amino acids could be counted with AA_spectra_number.py (https://github.com/MengTingHe2023/AA_spectra_number). CDR regions and suspicious sites were checked manually and shown by pLabel (v2.4.3.0).

### Antibodies Construction and Expression

The variable heavy chain (VH) and variable light chain (VL) gene sequences of anti-HA antibody were codon optimized and synthesized by RuiBiotech. The VH coding sequences of anti-HA were subsequently subcloned into a mouse immunoglobulin 2a (mIgG2a) heavy chain (HC) expression vector. Similarly, the VL coding sequence for anti-HA antibody was subcloned into a light chain expression vector. The heavy chain and light chain expression vectors were transfected into HEK293 cells at a weight ratio of 1:1. The HEK293 cells were cultured according to the manufacturer’s instructions (Thermo Fisher Scientific). The antibodies were subsequently purified using protein A beads (Smart-Lifesciences).

### Western Blot

10 ng or 5 ng of expressed proteins FLAG-GFP, FLAG-HA-GFP were spiked into 10 μg of total protein lysate from yeast cells, respectively. For Western blotting, 0.2 ng/mL of commercial or homemade antibodies was separately used as primary antibody, while 0.2 ng/mL of anti-mouse antibody served as the secondary antibody.

### Surface Plasmon Resonance (SPR) Analysis

The SPR analysis was conducted using a Biacore T200 instrument (Biacore, GE Healthcare). To determine the binding affinity of anti-HA antibodies to the HA peptide, the anti-HA antibodies (isotype mIgG2a) were captured with anti-mouse Fc antibodies (Biacore, GE Healthcare) which were immobilized on a CM5 sensor chip. Subsequently, the analytes, which consisted of a protein expressing the HA tag, were injected at various concentrations, in a serially diluted manner, into each flow cell. The Biacore T200 evaluation software was utilized to calculate the association rates (*Ka*), dissociation rates (*Kd*), and affinity constants (*K*^*D*^).

### Structure Prediction

We employed RosettaAntibody3^31^ for predicting the three-dimensional structure of both the HA peptide and its corresponding antibody based on their respective amino acid sequences. To generate models of HA-antibody complex, we performed computational docking using SnugDock^32^, resulting in a total of 1,000 models. Then we selected the HA-antibody complex model with the lowest score and compared Root Mean Square Deviation (RMSD) value with another antibody (PDB ID 5XCS). Subsequently, utilizing the Protein Local Optimization Program (PLOP)^33^ software, we refined the selected HA-antibody complexes by predicting the side chains of interface residues. Specifically, residues with any atoms within a 6 Å proximity to the HA peptide were targeted for optimization, including the HA peptide itself. All result data was analyzed by in-house analysis scripts.

### Illustration

Figure 1 was created using BioRender.

**Figure 1.**
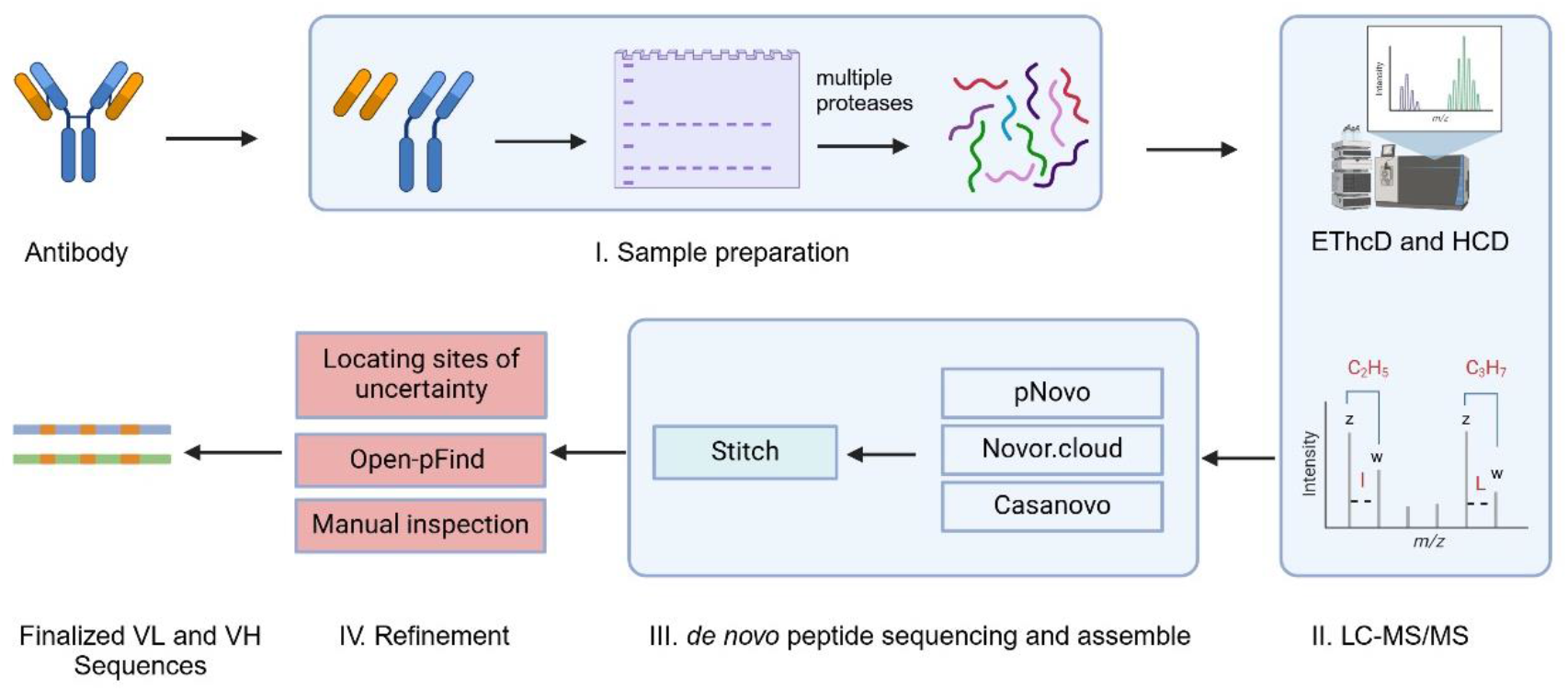
Overview of the workflow for *de novo* sequencing of monoclonal antibodies.

## RESULTS

### A workflow for *de novo* sequencing of a monoclonal antibody

The workflow described in this study consists of four steps: (1) Processing of a monoclonal antibody sample, (2) LC-MS/MS analysis, (3) *de novo* peptide sequencing and antibody sequence assembly, (4) sequence refinement. This workflow was developed using a homemade anti-FLAG monoclonal antibody as a reference sample. The sequence of this recombinant antibody is known, but we treated it as if it were not until the end of the workflow. After the workflow was established, we tested it on a commercial monoclonal antibody whose sequence is unknown to us. The details of the workflow are as follows.

#### (1) Sample processing

N-glycosylation is a common post-translational modification for antibodies, detected in the Fab region for 15-25% of the antibody species examined^34-36^. To preclude the possibility that glycosylation may interfere with peptide sequencing, the anti-FLAG antibody was deglycosylated with PNGase F. For comparison, another copy of the antibody sample was left untreated. To separate the light chain (LC) and the heavy chain (HC) from each other, we subjected the anti-FLAG antibody—deglycosylated or not—to reducing SDS-PAGE. Excised gel bands of the light chain and the heavy chain were reduced, alkylated, and digested in gel. Eight digestion conditions were set up to obtain overlapping peptides as much as possible, including Trypsin, Elastase, Pepsin, Chymotrypsin, Asp-N, GluC, Trypsin + AspN, Trypsin + GluC. The resulting peptides were extracted from gel bands for LC-MS/MS analysis.

#### (2) LC-MS/MS

High-resolution, accurate mass measurement is a prerequisite for *de novo* peptide sequencing in order to distinguish amino acid residues of similar masses, such as Q and K, or F and oxidized M. However, distinguishing I and L requires more since these two amino acids are of the same mass. Several fragmentation techniques have been developed to determine precisely whether a residue is I or L. These techniques include a low-energy collision-induced dissociation multistage mass analysis method, CID MSn, I unlike L, generates a 69 Da ion^37^, MS3 (ETD-HCD) approach^38^ and more recently, Electron Transfer Higher-energy Collision Dissociation (EThcD)^39, 40^. In the EThcD spectrum of a residue is I or L, it is observed that a z-ion with I or L at the very N-terminus is often accompanied by a w ion, which results from side-chain cleavage of the N-terminal I or L. In the case of I, an ethyl group is cleaved off, resulting in a mass difference of 29.039 Da (C_2_H_5_) between the z ion and its cognate w ion. In the case of L, an isopropyl group is cleaved off, resulting in a mass difference of 43.055 Da (C_3_H_7_) between the z ion and its cognate w-ion.

Taken the above into consideration, we chose to carry out LC-MS/MS analysis on a Q-Qrbitrap mass spectrometer with EThcD function. Each sample was analyzed twice to collect HCD spectra and EThcD spectra. The HCD and EThcD spectra could be collected in one run, but not all *de novo* software tools can process such data (e.g., Maxnovo), so we collected them separately. In brief, we analyzed each sample twice on a Fusion Lumos instrument over a 90-min reverse-phase LC run, once for HCD and once for EThcD.

#### (3) *de novo* peptide sequencing and antibody sequence assembly

We tested four publicly available, *de novo* sequencing software tools that are compatible with Stitch. They are pNovo, Casanovo, Novor.Cloud, and Maxnovo, all free of charge.

We began by inputting the *de novo* sequencing results of both the heavy chain and the light chain, HCD and EThcD data all included, into Stitch. One key parameter in Stitch is the cutoff value of Average Local Confidence (ALC). ALC indicates the confidence level of a peptide sequence assigned to an MS2 spectrum. As the ALC cutoff value increases, fewer but more reliable peptide sequences are retained by Stitch for subsequent antibody sequence assembly. At the default setting, Stitch failed to reconstruct the sequence of CDRH3, no matter which set of *de novo* sequence results were used as input (**Fig. S1)**. So, we optimized the ALC cutoff value for each of the *de novo* sequencing software used. The range of ALC values is 0-100 for pNovo, Maxnovo, and Novor.Cloud, or -1∼1 for Casanovo. We tested 5 or 6 ALC cutoff values from mid-point to the highest and found that at ALC cutoff ≥70, pNovo and Novor.Cloud, but not Maxnovo or Casanovo, could arrive at the accurate CDRH3 sequence.

Concerned that simultaneous assembly of the light chain and the heavy chain might bring an unnecessary burden to Stitch with Maxnovo and Casanovo sequences, we optimized the ALC cutoff value once again while keeping the assembly of LC and HC separate. As shown in **Fig. S3**, all except the Maxnovo results enabled reconstruction of the CDRH3 sequence at 100% accuracy through Stitch. For pNovo, Novor.Cloud, and Casanovo sequences, the recommended ALC cutoff values for LC/HC are, respectively, 60/70, 50/90, and 0.9/0.9. Deglycosylation hardly affected the optimization result of the ALC cutoff values **(Fig. S2-S5)**. At the recommended ALC cutoff settings with other parameters left at the default values, the reconstructed full-length LC and HC sequences are nearly 100% accurate. The occasional errors that remain can be classified into three types: ambiguity of isomers (I/L), ambiguity of same-mass amino acid groups (GG/N, GA/Q, etc.), and ambiguity of deamidation (N/D) **(Fig. 2)**.

**Figure 2.**
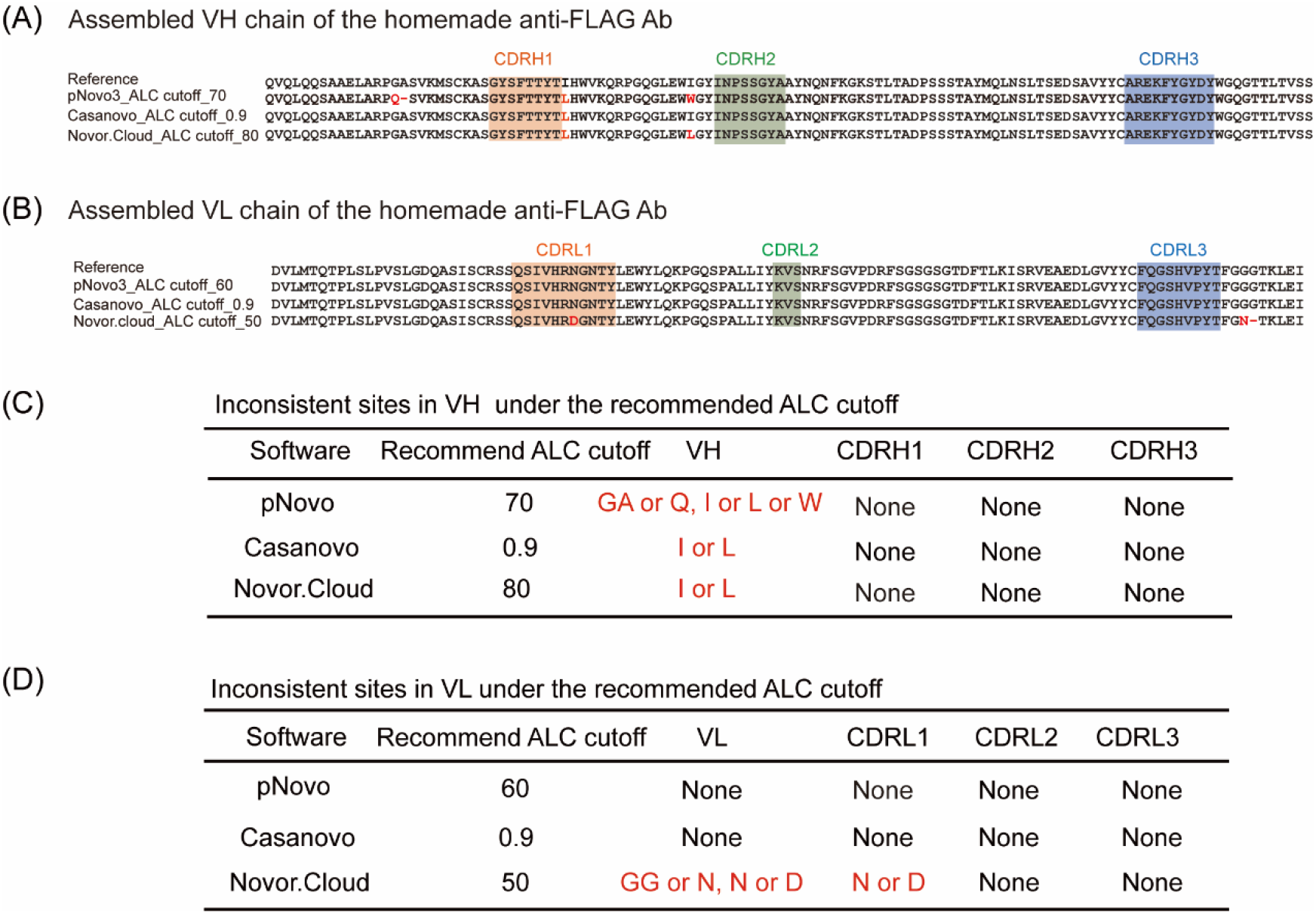
VH and VL sequences of the anti-FLAG antibody assembled under the recommended ALC cutoff values. (A, C) For VH, ≥98% sequence accuracy achieved, with the remaining inconsistent sites colored red. (B, D) For VL ≥98% sequence accuracy achieved, with the remaining inconsistent sites colored red.

#### (4) Sequence refinement

To rid the few remaining errors of the reconstructed antibody sequence, we developed a three-tiered sequence refinement strategy as follows.

First, sites of uncertainty were located by template and cross-reference alignment. Stitch takes a template-guided approach to assemble *de novo* sequence reads. It performs this task twice to reconstruct the sequence of a monoclonal antibody. The template sequence(s) used in the first round can be defined by users. In our case, the anti-FLAG antibody in question is a mouse IgG, and a multitude of mouse IgG antibody sequences are already included in the curated template sequence database in the installation package of Stitch, along with human antibody sequences, and rabbit antibody sequences, et,al. So we used the entire template sequence database of Stitch for the first assembly run. This resulted in a single template sequence for the second assembly run, leading to the final consensus sequence of the anti-FLAG antibody. These two sequences were compared to initiate the refinement procedure, which was helpful to locate questionable amino acid sites for further improvement **(Fig. 3A)**.

**Figure 3.**
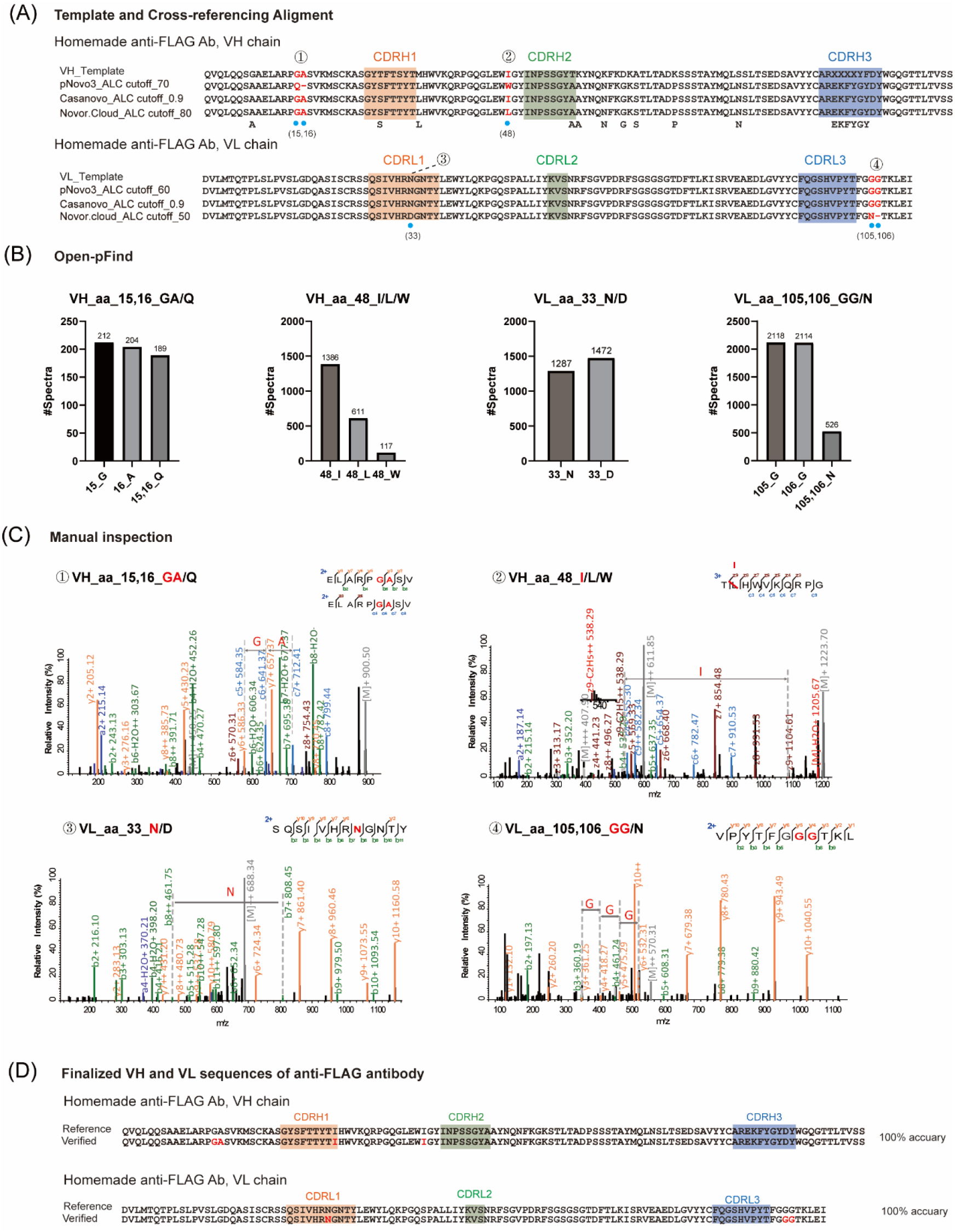
Procedure of sequence refinement. (A) Alignment of the best-scoring template sequence and the consensus sequences obtained from the pNovo, Casanovo, and Novor.Cloud *de novo* peptide sequence reads under the recommended ALC cutoff values. Red letters and blue dots mark the sites of ambiguity identified from comparison between consensus sequences and comparison between the template and consensus sequences, respectively. Single-letter amino acids below the alignment indicate correction of the template sequence by consensus sequences at the given position. (B) Open-pFind search results. The number of MS2 spectra supporting the specified identity of a given site is shown. (C) Manual inspection of MS2 to finalize the identity for each site of ambiguity. Representative MS2 spectra are shown. (D) Refined VH and VL sequences are identical to the reference sequences of the anti-FLAG antibody.

Next, we compared the antibody sequences reconstructed by Stitch based on the *de novo* sequencing reads from pNovo, Casanovo, and Novor.Cloud to minimize uncertain sites. For any given position, if the three consensus sequences do not agree with one another, then it is considered an ambiguous site. If they agree with one another but differ from the best scoring template in the second round of template matching, the amino acid identity given by the consensus sequences is taken as the final answer unless the discrepancy involves amino acid groups of the same mass (I/L, GG/N, AG/GA, etc.) or deamidated Q or N. In the latter case, the site in question is marked as an ambiguous site **(Fig. 3A)**.

Second, using the open search mode of pFind3, which offers an advantage of identifying point mutations, we resolved much of the ambiguity detected from above. Briefly, a protein sequence database was built out of the antibody sequences reconstructed by Stitch. The intermediate template sequences and the final consensus sequences assembled by Stitch out of the pNovo, Casanovo and Novor.Cloud sequence reads were all included. This database was then used to search the original HCD and EThcD spectra. As shown in **Fig. 3B and Fig. 3C**, three ambiguous sites GA/Q, N/D and GG/N were resolved by comparing spectral counts of the candidate sequences containing these sites, which led to GA, N, and GG as the final answers, respectively. Since pFind does not consider w-ions, it cannot differentiate between I and L.

Lastly, to all the questionable sites and all the CDR regions, we gave additional scrutiny by manually inspecting the MS2 spectra underlying the final sequence identity. In this way, the last questionable site I/L was also resolved **(Fig. 3C and Fig. S6)**. At the end of the three-tiered sequence refinement procedure as described above, we arrived at 100% sequence accuracy for the anti-FLAG antibody **(Fig. 3D)**.

### Application of the workflow to *de novo* sequencing of anti-HA Antibody (TANA2)

To test the effectiveness of our do-it-yourself *de novo* antibody sequencing workflow, we used it to decode an anti-HA antibody of unknown sequences. As shown in **Fig. 4A**, alignment of the best scoring template sequence with the three consensus sequences reconstructed by Stitch from pNovo, Casanovo, and Novor.Cloud *de novo* peptide sequence reads identified five ambiguous sites. The last of these sites, located less than 10 amino acids downstream of the CDR3 segment of the light chain is about isomeric ambiguity between I and L. The other four involve ambiguity of amino acids of different mass, and thus could be helped by database search if all the possibilities are included in the protein sequence database. We then conducted such database search using pFind3 under its open search mode. The Open-pFind search result resolved the ambiguity associated with the four sites **(Fig. 4B)**, as hundreds or thousands more MS2 spectra were assigned to one candidate versus the other. According to the pFind search results, HC (aa15,16) is GA nor Q, HC (aa99) has no additional serine residue LC (aa33) is N not D, LC (aa105, 106) is GG, not N. Lastly, we manually verified a number of highest-scoring MS2 spectra for each of the five ambiguous sites **(Fig. 4C)**. Through this effort, assignment of the last ambiguous sites was finalized: NC (aa111) is I, not L **(Fig. 4C)**. Manual inspection of MS2 was also conducted for the CDR regions **(Fig. S7)**. The light chain and heavy chain sequences obtained through this workflow are the same as the sequences determined by PEAKS AB **(Fig. 4D)**.

**Figure 4.**
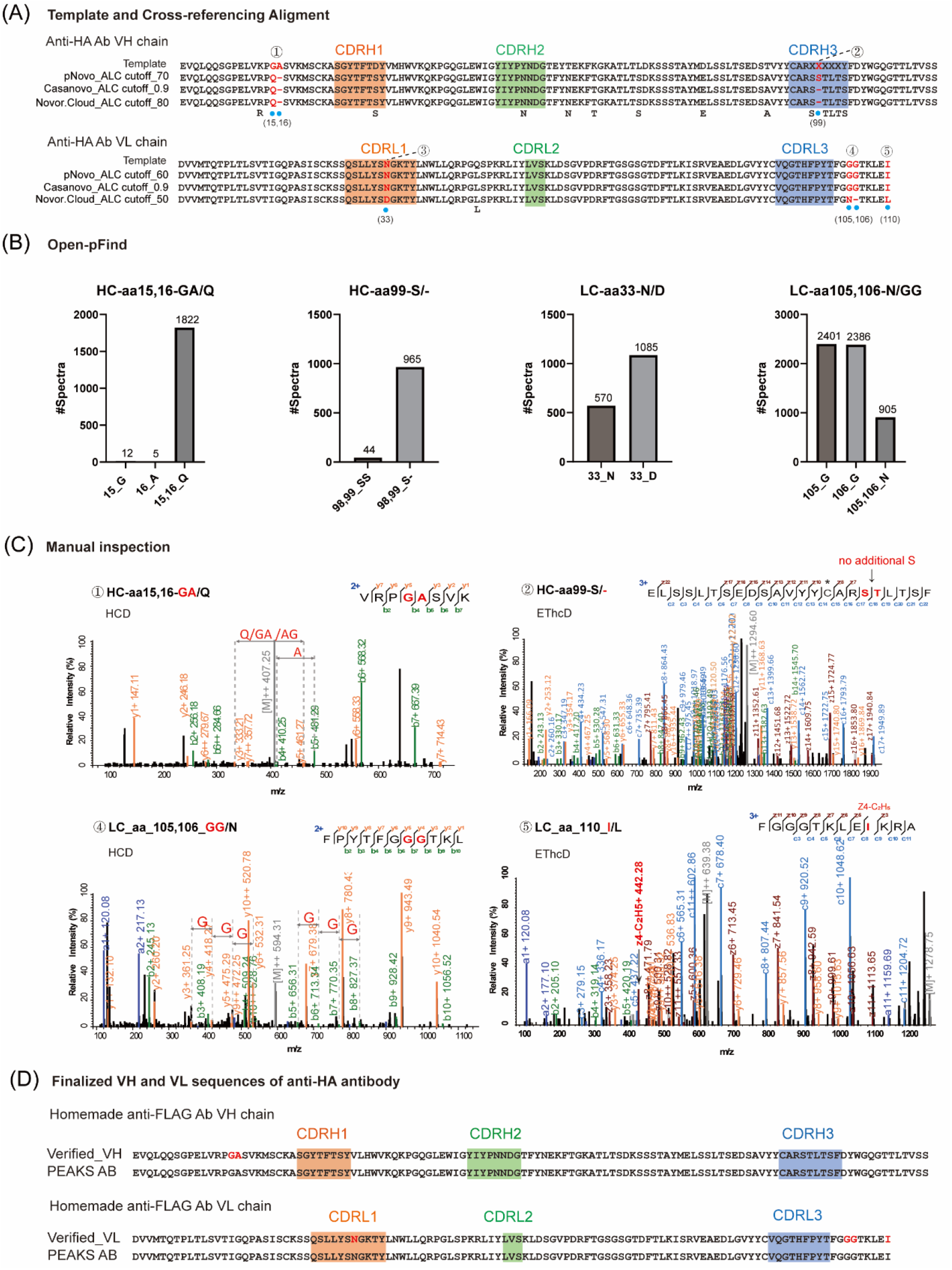
*De novo* sequencing of anti-HA antibody (TANA2) using the workflow. (A) Alignment of the best-scoring template sequence and the consensus sequences obtained from the pNovo, Casanovo, and Novor.Cloud *de novo* peptide sequence reads under the recommended ALC cutoff values. Red letters and blue dots mark the sites of ambiguity identified from comparison between consensus sequences and comparison between the template and consensus sequences, respectively.

Single-letter amino acids below the alignment indicate correction of the template sequence by consensus sequences at the given position. (B) Open-pFind search results. The number of MS2 spectra supporting the specified identity of a given site is shown. (C) Manual inspection of MS2 to finalize the identity for each site of ambiguity. representative MS2 spectra are shown. Carbamidomethylated C is labeled with * on the top in the peptide sequence displayed in the annotated MS2. (D) Refined VH and VL sequences of the anti-HA antibody.

### The specificity of homemade anti-HA antibody is as good as commercial version on western blot

To validate the sequences of the anti-HA antibody obtained as above, we had the DNA coding sequences of the variable regions of the light chain and heavy chain synthesized and subcloned into antibody expression vectors. The homemade anti-HA antibody was isolated from the culture medium of HEK293 cells co-transfected with the light and heavy chain expression vectors. As shown by western blotting **(Fig. 5)**, our DIY antibody and the commercial antibody performed equally well. Both recognized a FLAG-HA-GFP fusion protein (FHG), neither retained on the FLAG-GFP protein band (FG, negative control). When the fourth Asp residue of the HA epitope tag was mutated to Gly, the D4G variant of the FLAG-HA(D4G)-HA fusion protein was still recognizable for both versions of the anti-HA antibody, although the WB signal weakened pronouncedly

**Figure 5.**
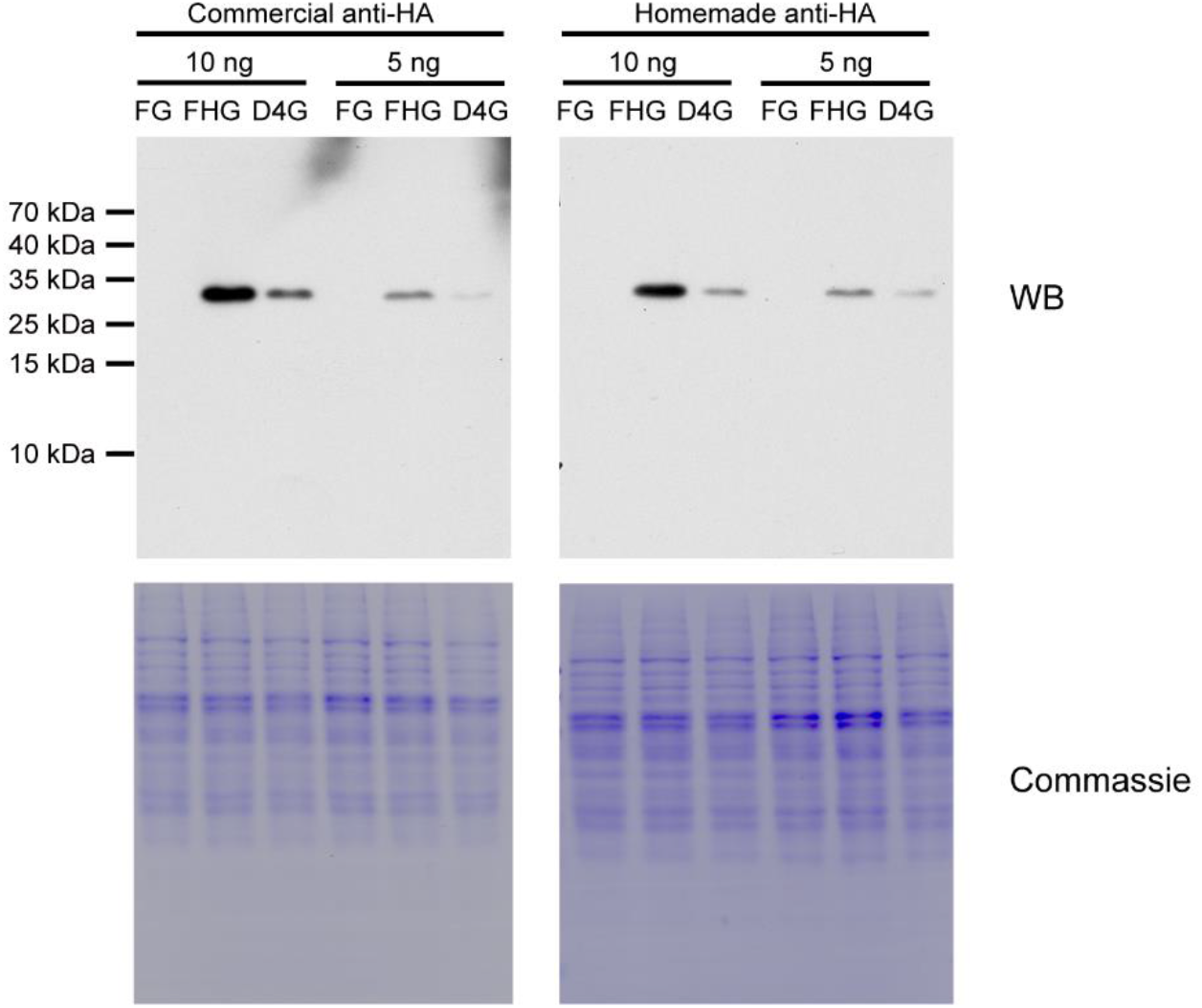
Western blot validation of the DIY antibody sequencing result. The resultant recombinant anti-HA antibody and the commercial one were compared side by side. 10 or 5 ng of purified FLAG-GFP (FG), FLAG-HA-GFP (FHG) or FLAG-HA (D4G)-GFP protein was mixed with 10 μg of background proteins from a yeast lysate, and separated on SDS-PAGE for WB. WB (above), Coomassie stain (below).

### Homemade anti-HA antibody exhibits similar affinity to the commercial version in binding with the HA-tagged protein

We further characterized the binding kinetics of the homemade and the commercial (TANA2) anti-HA antibodies. By Surface Plasmon Resonance (SPR) assay, the *K*_*a*_, *K*_*d*_ *and K*_*D*_ values are 7.25×10^-4^ M^-1^s^-1^, 2.13×10^-3^ s^-1^ and 2.94 ×10^-8^ M, respectively, for our DIY anti-HA antibody. These values are essentially the same as those for the commercial antibody, which are 8.70×10^-4^ M^-1^s^-1^, 2.58×10^-3^ s^-1^, and 2.97×10^-8^ M, respectively **(Fig. 6)**. Lastly, we used RosettaAntibody3 to model the 3D-structure of the antibody-antigen complex based on the anti-HA sequences obtained *de novo*. As shown **(Fig. S8)**, seven hydrogen bonds could form between the HA peptide and the antibody, and the H-bonded residue pairs are HA-Asp (4)-VH-Tyr (52), HA-Asp (4)-VH-Asn (53), HA-Val (5)-VH-Ser (31), HA-Pro (6)-VH-Thr (96), HA-Asp (7)-VH-Tyr (50), HA-Asp (7)-VL-Tyr (96), HA-Tyr (8)-VH-Leu (97).

**Figure 6.**
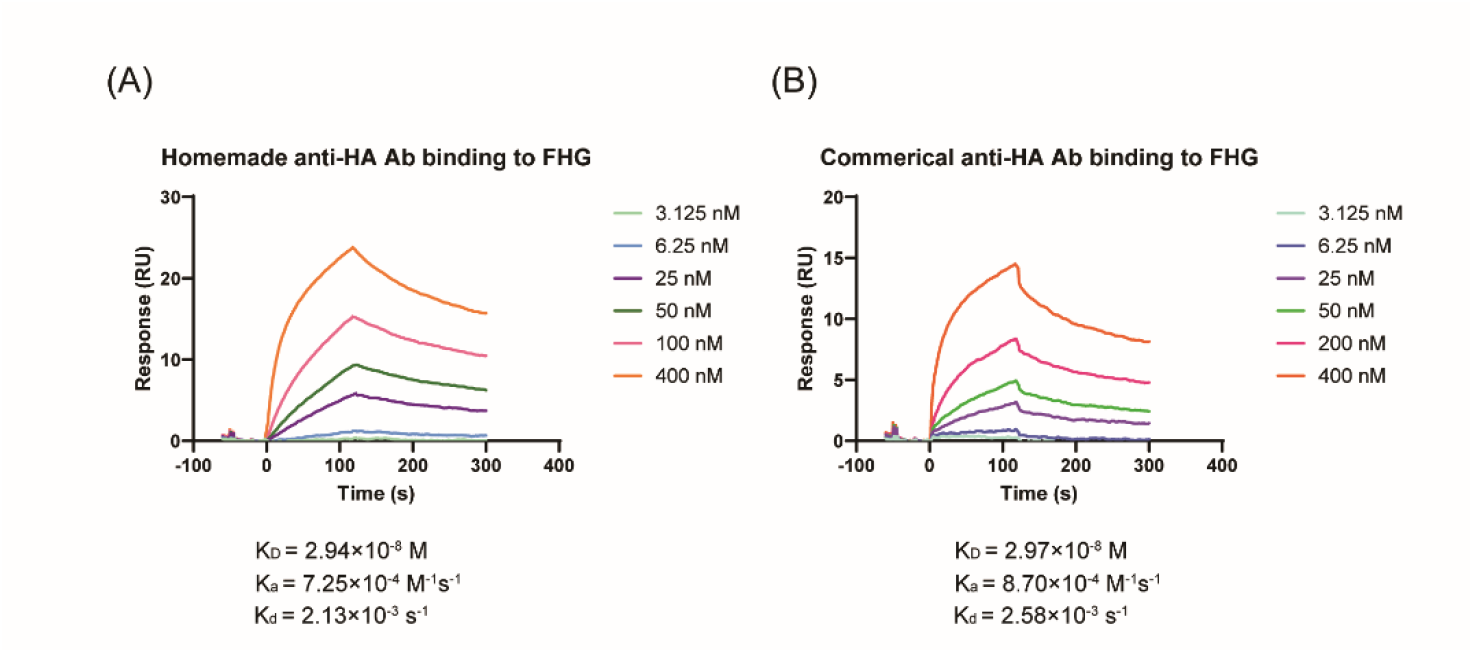
SPR measurement of binding kinetics of the DIY anti-HA antibody in comparison to the commercial antibody. Purified FLAG-HA-GFP was used at the indicated concentrations.

## DISCUSSION

In biomedical research, the need for *de novo* antibody sequencing is on the rise. MS-based *de novo* sequencing technology has advanced to a point that success is all but guaranteed for *de novo* sequencing of a purified monoclonal antibody by commercial service providers. Here, we provide a do-it-yourself workflow that could empower any MS facility or research laboratory to sequence monoclonal antibodies at will. This workflow provides 100% accurate VH and VL sequences at minimal cost, thanks to the free software tools including Stitch, pNovo, Casanovo and Novor.Cloud. We expect that this workflow will encourage increased participation from a broader range of MS laboratories to expand the sequence database of antibodies.

Although effective, MS-based *de novo* sequencing could benefit from further improvement in the following aspects. First, the Casanovo training model, Casanovo is currently trained on CID/HCD spectra of tryptic peptides and mixed non-tryptic peptides. However, to decode antibody sequences completely and accurately, it is necessary to incorporate peptides of distinct enzyme specificity, such as those generated by Asp-N, Glu-C, Chymotrypsin, etc. Casanovo would also improve if trained on EThcD spectra, which are of great value to *de novo* sequencing. Second, distinguishing between amino acids or amino acid combinations of identified mass values, e.g., GA/Q, GG/N, VA/GL, and more. This would require higher-quality MS2 data with fragment ions providing full coverage of the peptide backbone. Targeted MS approach and complementary fragmentation methods are possibilities to increase sequence coverage. Using deep learning technology to predict retention time and other physical-chemical properties of isobaric peptides is another way to resolve this issue, which is already employed by PEAKS AB. Third, distinguishing I and L. EThcD is effective in this regard, but sometimes the intensity if the relevant to the w- and z-ions are too low. Elevated production of the relevant w- and z-ions would help resolve the I/L ambiguity. C-terminal labeling with arginine methyl ester is recently reported to achieve such effect^41^. Lastly, on the software side, enhancing the identification accuracy for CDRH3 is most welcome.

## ASSOCIATED CONTENT

VL and VH sequences assembled separately for the anti-FLAG antibody, using Stitch in combination with pNovo, Novor.Cloud, Maxnovo, or Casanovo on default ALC cutoff values 9, 10, 0, 0 **(Fig. S1)**. VL and VH sequences aseembled together for the homemade anti-FLAG antibody using Stitch in combination with pNovo, Novor.Cloud, Maxnovo or Casanovo on different ALC cutoff values **(Fig. S2)**. VL and VH sequences assembled separately for the anti-FLAG antibody, using Stitch in combination with pNovo, Novor.Cloud, Maxnovo, or Casanovo on different ALC cutoff values **(Fig. S3)**. VL and VH sequences assembled together after deglycosylation for the anti-FLAG antibody, using Stitch in combination with pNovo, Novor.Cloud, Maxnovo, or Casanovo on different ALC cutoff values **(Fig. S4)**. VL and VH sequences assembled separately after deglycosylation for the anti-FLAG antibody, using Stitch in combination with pNovo, Novor.Cloud, Maxnovo, or Casanovo on different ALC cutoff values **(Fig. S5)**. Spectra supporting the reconstructed CDR sequences of the homemade anti-FLAG antibody **(Fig. S6)**. Spectra supporting the reconstructed CDR sequences of the anti-HA antibody (TANA2) **(Fig. S7)**. A structure model of the anti-HA Ab in complex with the HA peptide, computed using RosettaAntibody 3 **(Fig. S8)**. Mass spectrometry proteomics data have been deposited into the iProX https://www.iprox.cn/page/PSV023.html;?url=17349291806600bAg, Password: iMbr

## AUTHOR INFORMATION

### Authors

Meng-Ting He — College of Life Sciences, Beijing Normal University, 19 Xinjiekouwai Avenue,

Beijing, 100875, China; National Institute of Biological Sciences, Beijing 102206, China; Tsinghua Institute of Multidisciplinary Biomedical Research, Tsinghua University, Beijing 100084, China.

Ning Li — National Institute of Biological Sciences, Beijing 102206, China; Tsinghua Institute of Multidisciplinary Biomedical Research, Tsinghua University, Beijing 100084, China.

Jian-Hua Wang — Changping District, Beijing 102206, China.

Zhi-Zhong Wei — National Institute of Biological Sciences, Beijing 102206, China; Tsinghua Institute of Multidisciplinary Biomedical Research, Tsinghua University, Beijing 100084, China.

Jie Feng — National Institute of Biological Sciences, Beijing 102206, China; Tsinghua Institute of Multidisciplinary Biomedical Research, Tsinghua University, Beijing 100084, China.

Wen-Ting Li — Bioinformatics Solutions Inc., Waterloo, ON N2L 6J2, Canada.

Jian-Hua Sui — National Institute of Biological Sciences, Beijing 102206, China; Tsinghua Institute of Multidisciplinary Biomedical Research, Tsinghua University, Beijing 100084, China.

Niu Huang — National Institute of Biological Sciences, Beijing 102206, China. Tsinghua Institute of Multidisciplinary Biomedical Research, Tsinghua University, Beijing 100084, China.

### Author Contributions

Meng-Ting He and Meng-Qiu Dong designed the project. Meng-Ting He carried out the MS experiments. Meng-Ting He and Ning Li analyzed the MS data. Wen-Ting Li decoded anti-HA antibody (TANA2) using PEAKs AB. Zhi-Zhong Wei was responsible for the cloning and expression of the synthetic recombinant anti-HA antibody (TANA2). Zhi-Zhong Wei performed the SPR experiments. Meng-Ting He conducted the Western blotting experiments. Jie Feng predicted the structure of the HA tag and the anti-HA antibody complex. Meng-Qiu Dong, Jian-Hua Sui, and Niu Huang supervised the project. Meng-Qiu Dong and Meng-Ting He drafted the manuscript.

### Notes

The authors declare no competing financial interest.

## ACKNOWLEDGMENTS

The authors would like to acknowledge support from Stitch developer Douwe Schulte, who gave us much guidance on how to use the software. Thanks to pNovo developers Hao Chi, Jin-Yang Li, and Jia-Xiang Ding for their support on software upgrades. Thanks to Jian-Hua Wang for the initial guidance.

## REFERENCES

(1) Lu, R. M.; Hwang, Y. C.; Liu, I. J.; Lee, C. C.; Tsai, H. Z.; Li, H. J.; Wu, H. C. Development of therapeutic antibodies for the treatment of diseases. J. Biomed. Sci. 2020, 27 (1), 1. DOI: 10.1186/s12929-019-0592-z.

(2) Ecker, D. M.; Jones, S. D.; Levine, H. L. The therapeutic monoclonal antibody market. Mabs 2015, 7 (1), 9–14. DOI: 10.4161/19420862.2015.989042.

(3) Gray, A.; Bradbury, A. R. M.; Knappik, A.; Plückthun, A.; Borrebaeck, C. A. K.; Dübel, S. Animal-free alternatives and the antibody iceberg. Nature Biotechnology 2020, 38 (11), 1234–1239. DOI: 10.1038/s41587-020-0687-9.

(4) Bradbury, A.; Plückthun, A. Reproducibility: Standardize antibodies used in research. Nature 2015, 518 (7537), 27–29. DOI: 10.1038/518027a.

(5) Azevedo Reis Teixeira, A.; Erasmus, M. F.; D’Angelo, S.; Naranjo, L.; Ferrara, F.; Leal-Lopes, C.; Durrant, O.; Galmiche, C.; Morelli, A.; Scott-Tucker, A.; et al. Drug-like antibodies with high affinity, diversity and developability directly from next-generation antibody libraries. MAbs 2021, 13 (1), 1980942. DOI: 10.1080/19420862.2021.1980942.

(6) Chiu, M. L.; Goulet, D. R.; Teplyakov, A.; Gilliland, G. L. Antibody Structure and Function: The Basis for Engineering Therapeutics. Antibodies (Basel) 2019, 8 (4). DOI: 10.3390/antib8040055.

(7) Almagro, J. C.; Fransson, J. Humanization of antibodies. Front Biosci 2008, 13 (5), 1619–1633. DOI: 10.2741/2786.

(8) Subas Satish, H. P.; Zeglinski, K.; Uren, R. T.; Nutt, S. L.; Ritchie, M. E.; Gouil, Q.; Kluck, R. M. NAb-seq: an accurate, rapid, and cost-effective method for antibody long-read sequencing in hybridoma cell lines and single B cells. MAbs 2022, 14 (1), 2106621. DOI: 10.1080/19420862.2022.2106621.

(9) Meyer, L.; López, T.; Espinosa, R.; Arias, C. F.; Vollmers, C.; DuBois, R. M. A simplified workflow for monoclonal antibody sequencing. PLoS One 2019, 14 (6), e0218717. DOI: 10.1371/journal.pone.0218717.

(10) Ma, B.; Zhang, K.; Hendrie, C.; Liang, C.; Li, M.; Doherty-Kirby, A.; Lajoie, G. PEAKS: powerful software for peptide de novo sequencing by tandem mass spectrometry. Rapid Commun. Mass Spectrom. 2003, 17 (20), 2337–2342. DOI: 10.1002/rcm.1196.

(11) Frank, A.; Pevzner, P. PepNovo: de novo peptide sequencing via probabilistic network modeling. Anal. Chem. 2005, 77 (4), 964–973. DOI: 10.1021/ac048788h.

(12) Chi, H.; Sun, R. X.; Yang, B.; Song, C. Q.; Wang, L. H.; Liu, C.; Fu, Y.; Yuan, Z. F.; Wang, H. P.; He, S. M.; et al. pNovo: de novo peptide sequencing and identification using HCD spectra. J Proteome Res 2010, 9 (5), 2713–2724. DOI: 10.1021/pr100182k.

(13) Ma, B. Novor: real-time peptide de novo sequencing software. J Am Soc Mass Spectrom 2015, 26 (11), 1885–1894. DOI: 10.1007/s13361-015-1204-0.

(14) Tran, N. H.; Zhang, X.; Xin, L.; Shan, B.; Li, M. De novo peptide sequencing by deep learning. PNAS 2017, 114 (31), 8247–8252. DOI: Proc Natl Acad Sci U S A10.1073/pnas.1705691114.

(15) Karunratanakul, K.; Tang, H. Y.; Speicher, D. W.; Chuangsuwanich, E.; Sriswasdi, S. Uncovering Thousands of New Peptides with Sequence-Mask-Search Hybrid De Novo Peptide Sequencing Framework. Mol Cell Proteomics 2019, 18 (12), 2478–2491. DOI: 10.1074/mcp.TIR119.001656.

(16) Gutenbrunner, P.; Kyriakidou, P.; Welker, F.; Cox, J. Spectrum graph-based de-novo sequencing algorithm MaxNovo achieves high peptide identification rates in collisional dissociation MS/MS spectra. 2021. DOI: 10.1101/2021.09.04.458985.

(17) Xu, X.; Yang, C.; He, Q.; Shu, K.; Xinpu, Y.; Chen, Z.; Zhu, Y.; Chen, T. PGPointNovo: an efficient neural network-based tool for parallel de novo peptide sequencing. Bioinform Adv 2023, 3 (1), vbad057. DOI: 10.1093/bioadv/vbad057 (acccessed 12/5/2023).

(18) Klaproth-Andrade, D.; Hingerl, J.; Smith, N. H.; Träuble, J.; Wilhelm, M.; Gagneur, J. Deep learning-driven fragment ion series classification enables highly precise and sensitive de novo peptide sequencing. 2023. DOI: 10.1101/2023.01.05.522752.

(19) Wu, S.; Luan, Z.; Fu, Z.; Wang, Q.; Guo, T. BiATNovo: A Self-Attention based Bidirectional Peptide Sequencing Method. 2023. DOI: 10.1101/2023.05.11.540352.

(20) Yang, T.; Ling, T.; Sun, B.; Liang, Z.; Xu, F.; Huang, X.; Xie, L.; He, Y.; Li, L.; He, F.; et al. Introducing pi-HelixNovo for practical large-scale de novo peptide sequencing. Brief Bioinform 2024, 25 (2). DOI: 10.1093/bib/bbae021.

(21) Guthals, A.; Clauser, K. R.; Bandeira, N. Shotgun protein sequencing with meta-contig assembly. Mol Cell Proteomics 2012, 11 (10), 1084–1096. DOI: 10.1074/mcp.M111.015768.

(22) Savidor, A.; Barzilay, R.; Elinger, D.; Yarden, Y.; Lindzen, M.; Gabashvili, A.; Adiv Tal, O.; Levin, Y. Database-independent Protein Sequencing (DiPS) Enables Full-length de Novo Protein and Antibody Sequence Determination. Mol Cell Proteomics 2017, 16 (6), 1151–1161. DOI: 10.1074/mcp.O116.065417.

(23) Mai, Z. B.; Zhou, Z. H.; He, Q. Y.; Zhang, G. Highly Robust de Novo Full-Length Protein Sequencing. Anal. Chem. 2022, 94 (8), 3467–3475. DOI: 10.1021/acs.analchem.1c03718.

(24) Schulte, D.; Peng, W.; Snijder, J. Template-Based Assembly of Proteomic Short Reads For De Novo Antibody Sequencing and Repertoire Profiling. Anal. Chem. 2022, 94 (29), 10391–10399. DOI: 10.1021/acs.analchem.2c01300.

(25) Schulte, D.; Snijder, J. A Handle on Mass Coincidence Errors in De Novo Sequencing of Antibodies by Bottom-up Proteomics. J Proteome Res 2024. DOI: 10.1021/acs.jproteome.4c00188.

(26) Tran, N. H.; Rahman, M. Z.; He, L.; Xin, L.; Shan, B.; Li, M. Complete De Novo Assembly of Monoclonal Antibody Sequences. Scientific reports 2016, 6 (1), 31730. DOI: 10.1038/srep31730.

(27) Bern, M.; Kil, Y. J.; Becker, C. Byonic: advanced peptide and protein identification software. Current protocols in bioinformatics 2012, Chapter 13, 13 20 11–13 20 14. DOI: 10.1002/0471250953.bi1320s40.

(28) Beslic, D.; Tscheuschner, G.; Renard, B. Y.; Weller, M. G.; Muth, T. Comprehensive evaluation of peptide de novo sequencing tools for monoclonal antibody assembly. Brief Bioinform 2023, 24 (1). DOI: 10.1093/bib/bbac542.

(29) Jian-Hua, W.; Zhou, G.; Xu, D.; Shu-Qun, L.; Yu-Liang, T.; Xiaoguang, L.; Chun, T.; Meng-Qiu, D. Fast cross-linking by DOPA2 promotes the capturing of a stereospecific protein complex over nonspecific encounter complexes. Biophysics Reports 2022, 8 (5-6), 239–252. DOI: 10.52601/bpr.2022.220014.

(30) Shao, G.; Cao, Y.; Chen, Z.; Liu, C.; Li, S.; Chi, H.; Dong, M. Q. How to use open-pFind in deep proteomics data analysis?-A protocol for rigorous identification and quantitation of peptides and proteins from mass spectrometry data. Biophys Rep 2021, 7 (3), 207–226. DOI: 10.52601/bpr.2021.210004.

(31) Weitzner, B. D.; Kuroda, D.; Marze, N.; Xu, J.; Gray, J. J. Blind prediction performance of RosettaAntibody 3.0: grafting, relaxation, kinematic loop modeling, and full CDR optimization. Proteins 2014, 82 (8), 1611–1623. DOI: 10.1002/prot.24534.

(32) Sircar, A.; Gray, J. J. SnugDock: paratope structural optimization during antibody-antigen docking compensates for errors in antibody homology models. PLoS Comput Biol 2010, 6 (1), e1000644. DOI: 10.1371/journal.pcbi.1000644.

(33) Jacobson, M. P.; Kaminski, G. A.; Friesner, R. A.; Rapp, C. S. Force Field Validation Using Protein Side Chain Prediction. The Journal of Physical Chemistry B 2002, 106 (44), 11673–11680. DOI: 10.1021/jp021564n.

(34) Holland, M.; Yagi, H.; Takahashi, N.; Kato, K.; Savage, C. O.; Goodall, D. M.; Jefferis, R. Differential glycosylation of polyclonal IgG, IgG-Fc and IgG-Fab isolated from the sera of patients with ANCA-associated systemic vasculitis. Biochim Biophys Acta 2006, 1760 (4), 669–677. DOI: 10.1016/j.bbagen.2005.11.021.

(35) Arnold, J. N.; Wormald, M. R.; Sim, R. B.; Rudd, P. M.; Dwek, R. A. The impact of glycosylation on the biological function and structure of human immunoglobulins. Annu. Rev. Immunol. 2007, 25, 21–50. DOI: 10.1146/annurev.immunol.25.022106.141702.

(36) Liu, S.; Liu, X. Chapter One - IgG N-glycans. In Advances in Clinical Chemistry, Makowski, G. S. Ed.; Vol. 105; Elsevier, 2021; pp 1–47.

(37) Armirotti, A.; Millo, E.; Damonte, G. How to discriminate between leucine and isoleucine by low energy ESI-TRAP MSn. J Am Soc Mass Spectrom 2007, 18 (1), 57–63. DOI: 10.1016/j.jasms.2006.08.011.

(38) Lebedev, A. T.; Damoc, E.; Makarov, A. A.; Samgina, T. Y. Discrimination of Leucine and Isoleucine in Peptides Sequencing with Orbitrap Fusion Mass Spectrometer. Analytical Chemistry 2014, 86 (14), 7017–7022. DOI: 10.1021/ac501200h.

(39) Samgina, T. Y.; Kovalev, S. V.; Tolpina, M. D.; Trebse, P.; Torkar, G.; Lebedev, A. T. EThcD Discrimination of Isomeric Leucine/Isoleucine Residues in Sequencing of the Intact Skin Frog Peptides with Intramolecular Disulfide Bond. J Am Soc Mass Spectrom 2018, 29 (5), 842–852. DOI: 10.1007/s13361-017-1857-y.

(40) Zhokhov, S. S.; Kovalyov, S. V.; Samgina, T. Y.; Lebedev, A. T. An EThcD-Based Method for Discrimination of Leucine and Isoleucine Residues in Tryptic Peptides. J Am Soc Mass Spectrom 2017, 28 (8), 1600–1611. DOI: 10.1007/s13361-017-1674-3.

(41) Le Bihan, T.; McDonald, Z.; Celejewski, K. R.; Liu, Q.; Ma, B. Enhancing De Novo Protein Sequencing through the C-Terminal Labeling Strategy: Resolving Isobaric Ambiguities by Electron-Transfer/Higher Energy Collision Dissociation (EThcD). Analytical Chemistry 2024. DOI: 10.1021/acs.analchem.4c03459.

