## Supporting Information for "Do-it-yourself *de novo* antibody sequencing workflow that achieves complete accuracy of the variable regions"

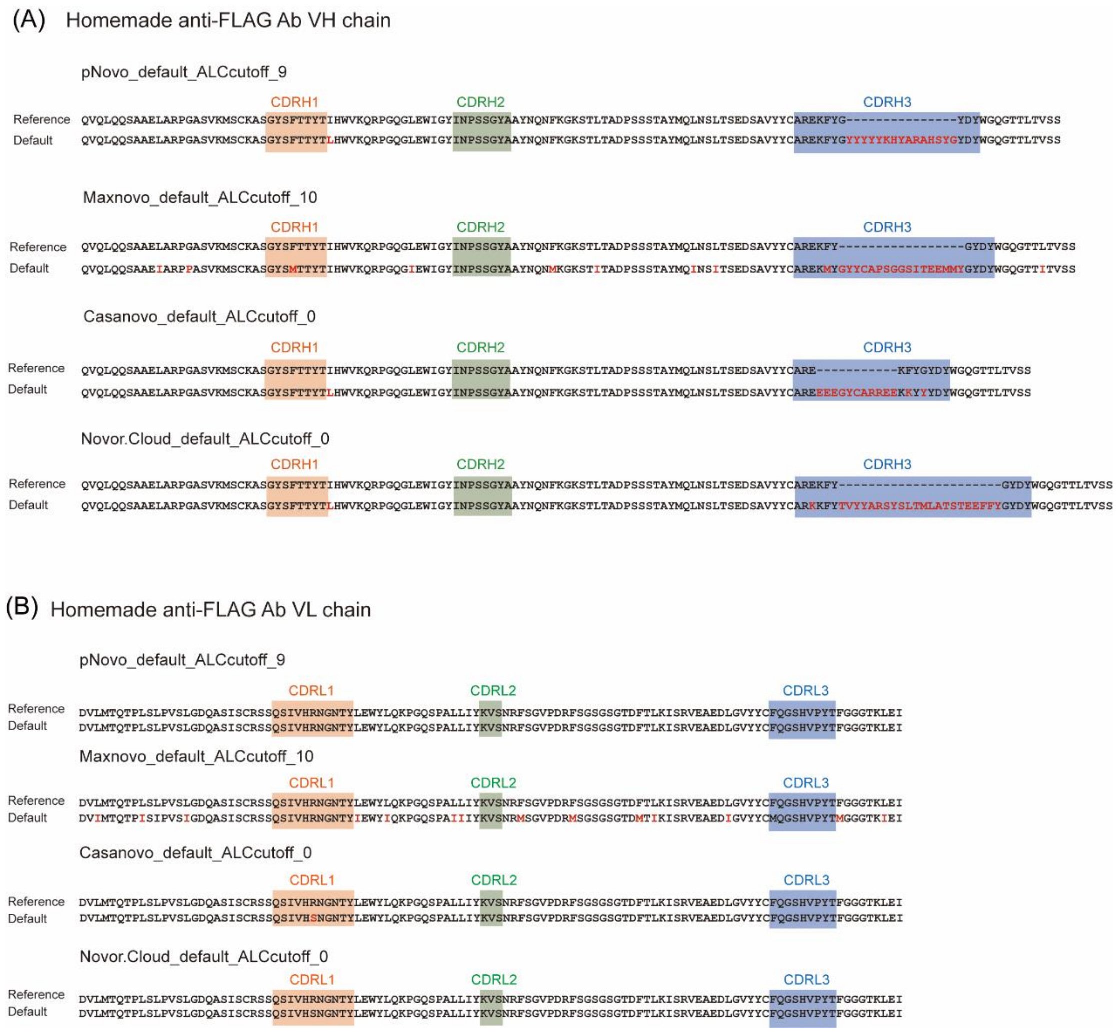


**Supplementary Figure 1.** VL and VH sequences assembled separately for the anti-FLAG antibody, using Stitch in combination with pNovo, Novor.cloud, Maxnovo, or Casanovo on default ALC cutoff values 9, 10, 0, 0. (A) All combinations failed to reconstruct CDRH3 on the default ALC cutoff. (B) Successful reconstruction of the VL sequence on the default ALC cutoff.

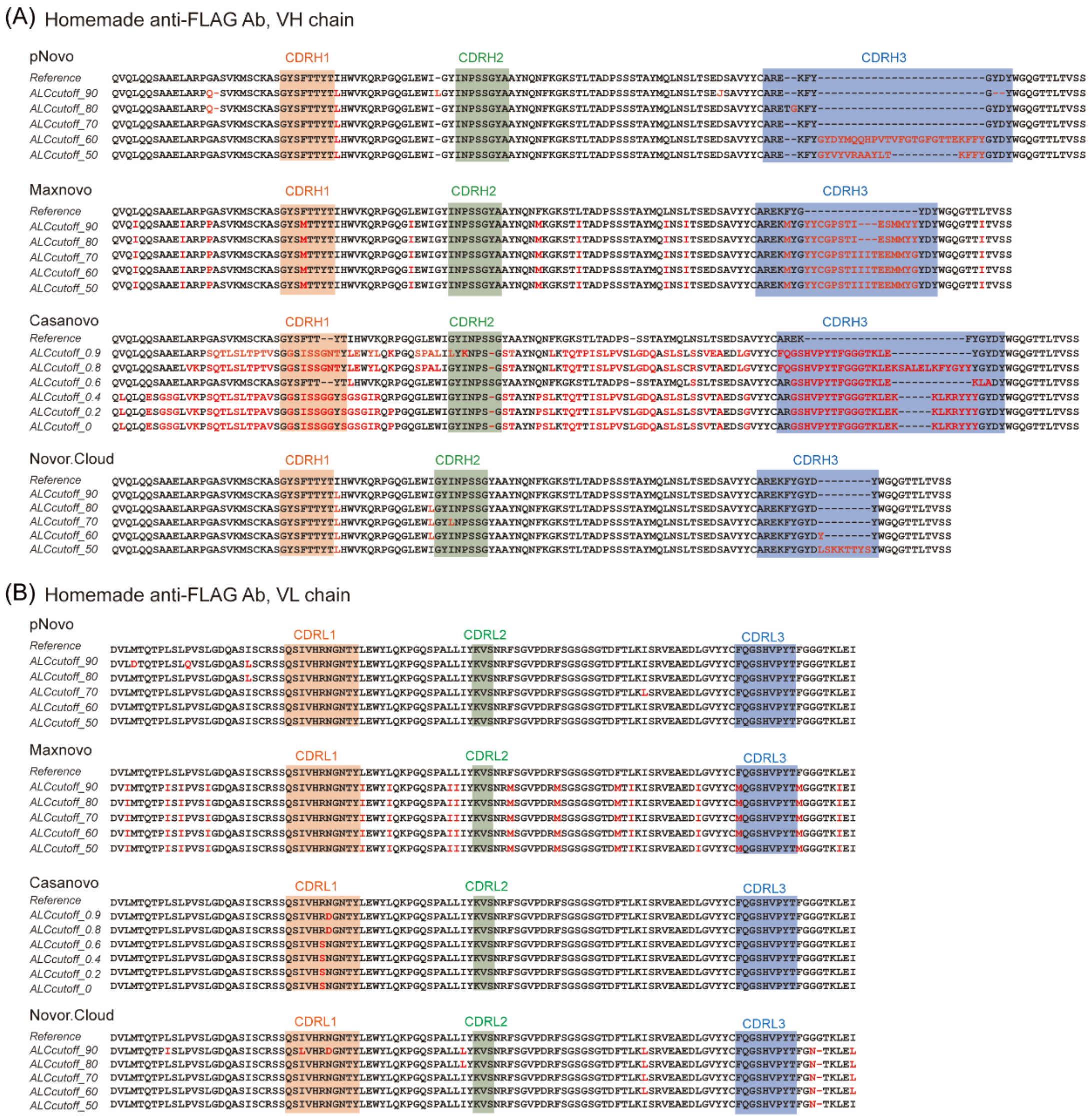


(C) Inconsistent sites in VH under the recommended ALC cutoff

| Software | Recommended ALC cutoff | VH | CDRH1 | CDRH2 | CDRH3 |
|---|---|---|---|---|---|
| pNovo | 70 | I or L | none | none | none |
| Novor.Cloud | 80 | I or L | none | none | none |

(D) Inconsistent sites in VL under the recommended ALC cutoff

| Software | Recommended ALC cutoff | VL | CDRL1 | CDRL2 | CDRL3 |
|---|---|---|---|---|---|
| pNovo | 60 | none | none | none | none |
| Casanovo | 0.9 | N or D | N or D | none | none |
| Novor.Cloud | 50 | GG or N | none | none | none |

**Supplementary Figure 2.** VL and VH sequences aseembled together for the homemade anti-FLAG antibody using Stitch in combination with pNovo, Novor.cloud, Maxnovo or Casanovo on different ALC cutoff values. (A, B) Reconstructed sequences of VH (A) and VL (B) by Stitch from *de novo* peptide sequence reads of pNovo, Maxnovo, Casanovo, or Novor.Cloud under different ALC cutoff values. Amino acids differ from the reference sequence are highlighted in red. (C, D) Recommended ALC cutoff values for VH (C) and VL (D) are not the same, although VH and VL are assembled together. No recommendation was made if sequence accuracy was below 98%. The remaining sequence deviations are listed in red letters.

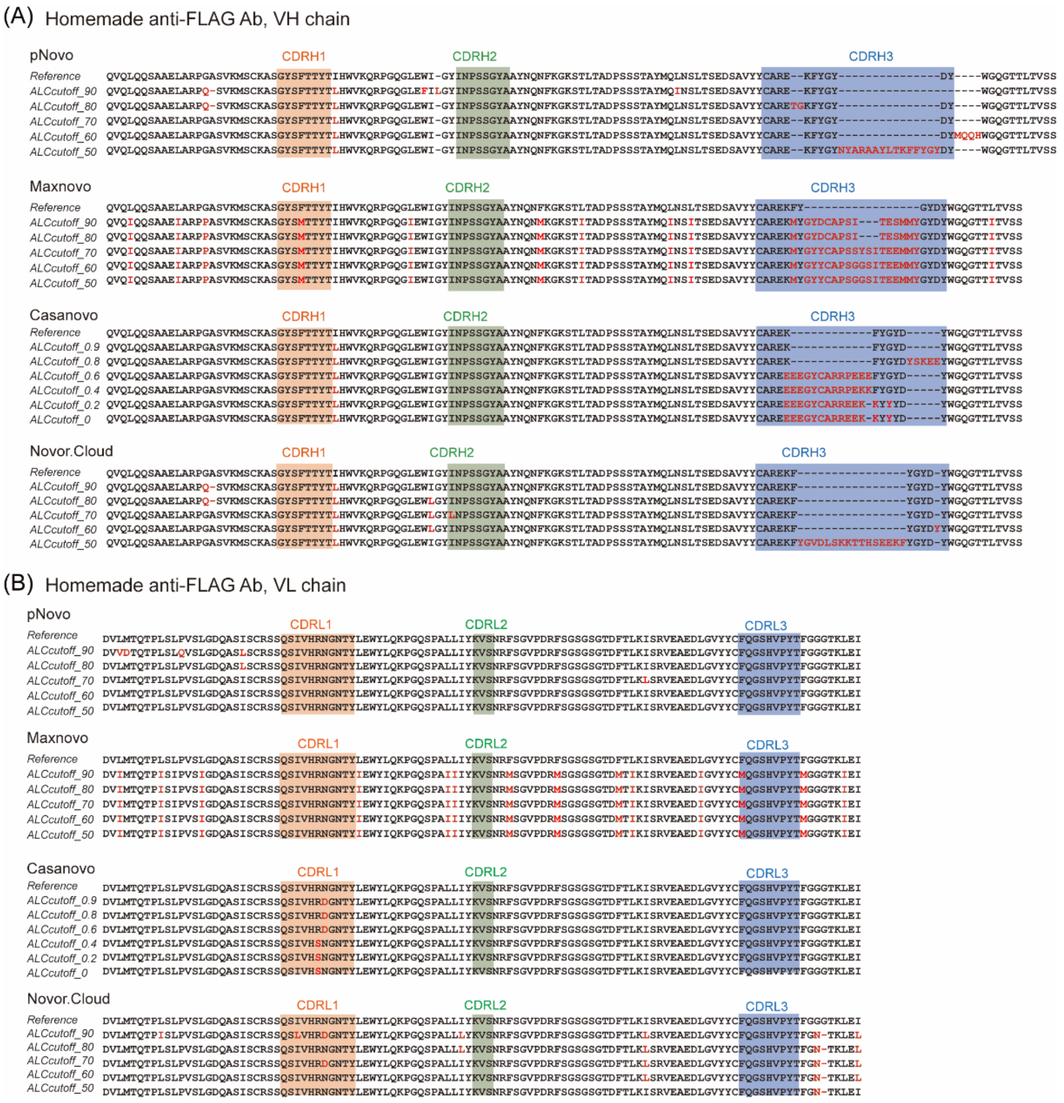


(C) Inconsistent sites in VH under the recommended ALC cutoff

| Software | Recommend ALC cutoff | VH | CDRH1 | CDRH2 | CDRH3 |
|---|---|---|---|---|---|
| pNovo | 70 | I or L | none | none | none |
| Casanovo | 0.9 | I or L | none | none | none |
| Novor.Cloud | 80 | GA or Q, I or L | none | none | none |

(D) Inconsistent sites in VL under the recommended ALC cutoff

| Software | Recommend ALC cutoff | VL | CDRL1 | CDRL2 | CDRL3 |
|---|---|---|---|---|---|
| pNovo | 60 | none | none | none | none |
| Casanovo | 0.9 | N or D | N or D | none | none |
| Novor.Cloud | 50 | GG or N | none | none | none |

**Supplementary Figure 3.** VL and VH sequences assembled separately for the anti-FLAG antibody, using Stitch in combination with pNovo, Novor.cloud, Maxnovo, or Casanovo on different ALC cutoff values. (A, B) Reconstructed sequences of VH (A) and VL (B) by Stitch from *de novo* peptide sequence reads of pNovo, Maxnovo, Casanovo, or Novor.Cloud under different ALC cutoff values. Amino acids differ from the reference sequence are highlighted in red. (C, D) Recommended ALC cutoff values for VH (C) and VL (D) are not the same. No recommendation was made if sequence accuracy was below 98%. The remaining sequence deviations are listed in red letters.

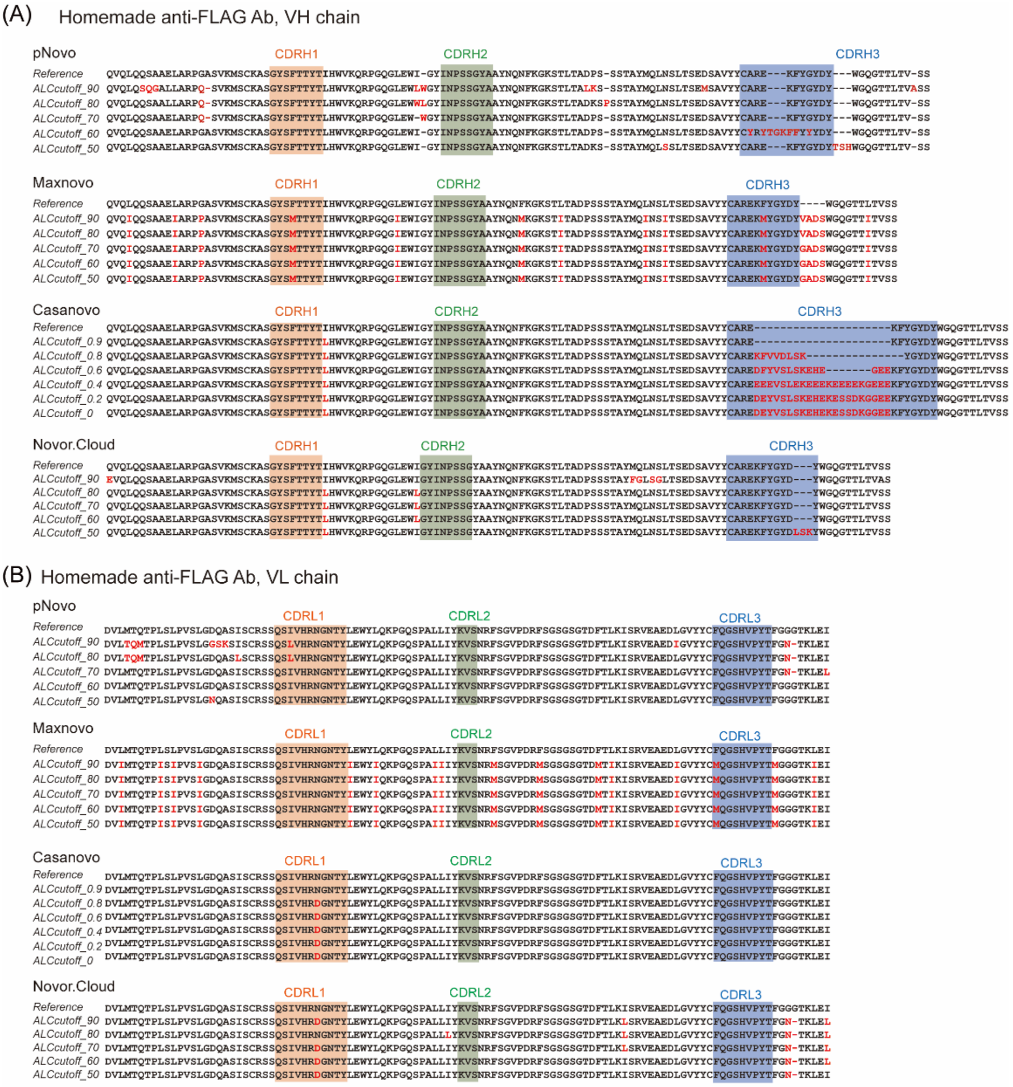


(C) Inconsistent sites in VH under the recommended ALC cutoff

| Software | Recommend ALC cutoff | VH | CDRH1 | CDRH2 | CDRH3 |
|---|---|---|---|---|---|
| pNovo | 70 | GA or Q, I or L or W | none | none | none |
| Casanovo | 0.9 | I or L | none | none | none |
| Novor.Cloud | 80 | I or L | none | none | none |

(D) Inconsistent sites in VL under the recommended ALC cutoff

| Software | Recommend ALC cutoff | VL | CDRL1 | CDRL2 | CDRL3 |
|---|---|---|---|---|---|
| pNovo | 60 | none | none | none | none |
| Casanovo | 0.9 | none | none | none | none |
| Novor.Cloud | 50 | GG or N, N or D | N or D | none | none |

**Supplementary Figure 4.** VL and VH sequences assembled together after deglycosylation for the anti-FLAG antibody, using Stitch in combination with pNovo, Novor.Cloud, Maxnovo, or Casanovo on different ALC cutoff values. (A, B) Reconstructed sequences of VH (A) and VL (B) by Stitch from *de novo* peptide sequence reads of pNovo, Maxnovo, Casanovo, or Novor.Cloud under different ALC cutoff values. Amino acids differ from the reference sequence are highlighted in red. (C, D) Recommended ALC cutoff values for VH (C) and VL (D) are not the same, although VH and VL are assembled together. No recommendation was made if sequence accuracy was below 98%. The remaining sequence deviations are listed in red letters.

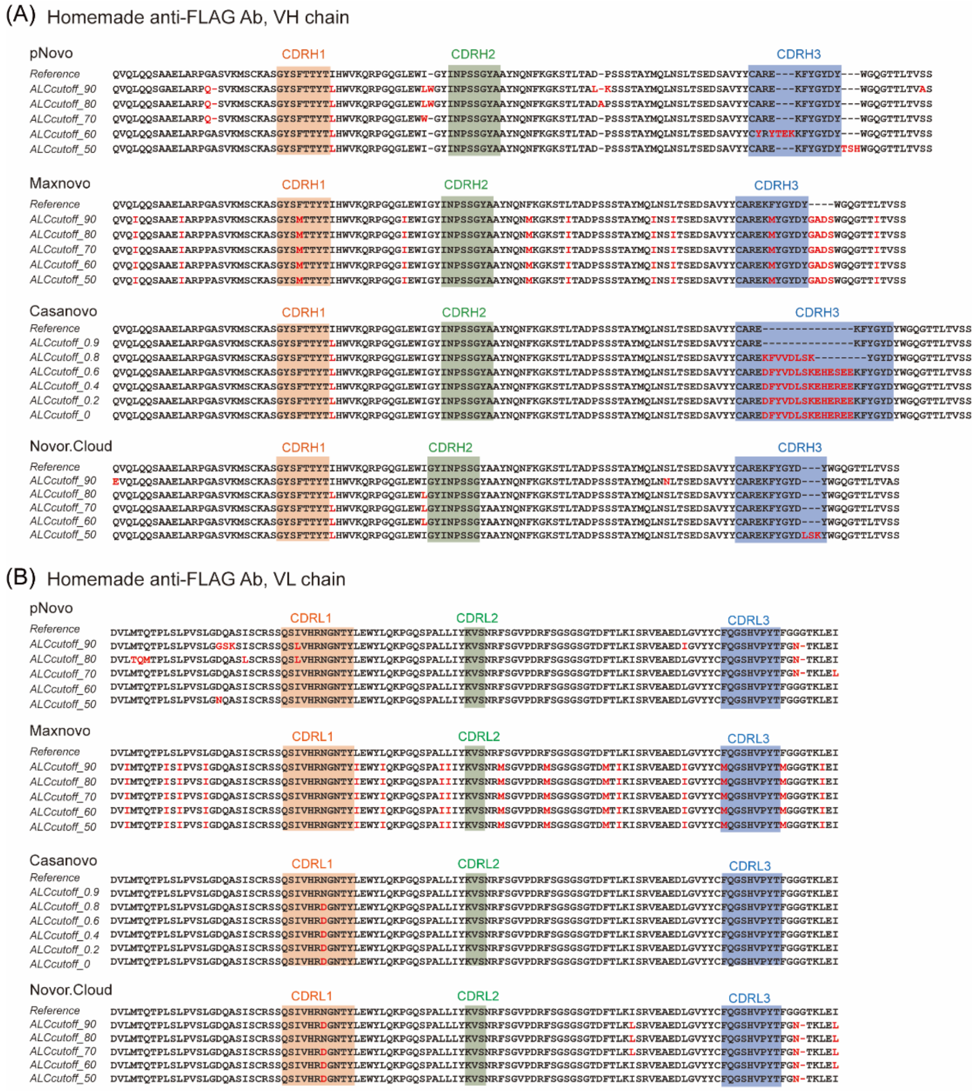


(C) Inconsistent sites in VH under the recommended ALC cutoff

| Software | Recommend ALC cutoff | VH | CDRH1 | CDRH2 | CDRH3 |
|---|---|---|---|---|---|
| pNovo | 70 | GA or Q, I or L or W | none | none | none |
| Casanovo | 0.9 | I or L | none | none | none |
| Novor.Cloud | 80 | I or L | none | none | none |

(D) Inconsistent sites in VL under the recommended ALC cutoff

| Software | Recommend ALC cutoff | VL | CDRL1 | CDRL2 | CDRL3 |
|---|---|---|---|---|---|
| pNovo | 60 | none | none | none | none |
| Casanovo | 0.9 | none | none | none | none |
| Novor.Cloud | 50 | GG or N, N or D | N or D | none | none |

**Supplementary Figure 5.** VL and VH sequences assembled separately after deglycosylation for the anti-FLAG antibody, using Stitch in combination with pNovo, Novor.Cloud, Maxnovo, or Casanovo on different ALC cutoff values. (A, B) Reconstructed sequences of VH (A) and VL (B) by Stitch from *de novo* peptide sequence reads of pNovo, Maxnovo, Casanovo, or Novor.Cloud under different ALC cutoff values. Amino acids differ from the reference sequence are highlighted in red. (C, D) Recommended ALC cutoff values for VH (C) and VL (D) are not the same. No recommendation was made if sequence accuracy was below 98%. The remaining sequence deviations are listed in red letters.

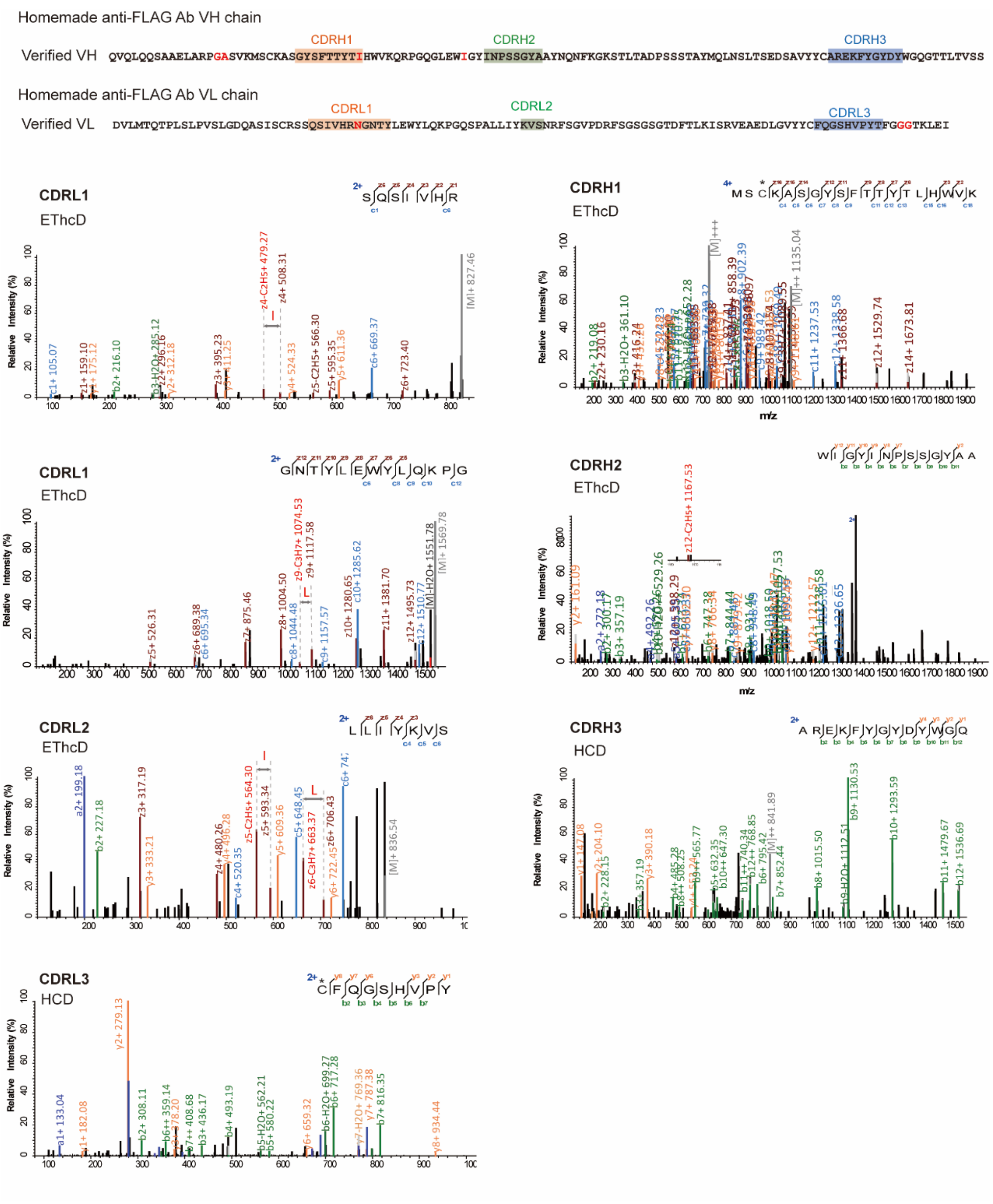


**Supplementary Figure 6.** Spectra supporting the reconstructed CDR sequences of the homemade anti-FLAG antibody. Carbamidomethylated C is labeled with * on the top in the peptide sequence displayed in the annotated MS2 spectra.

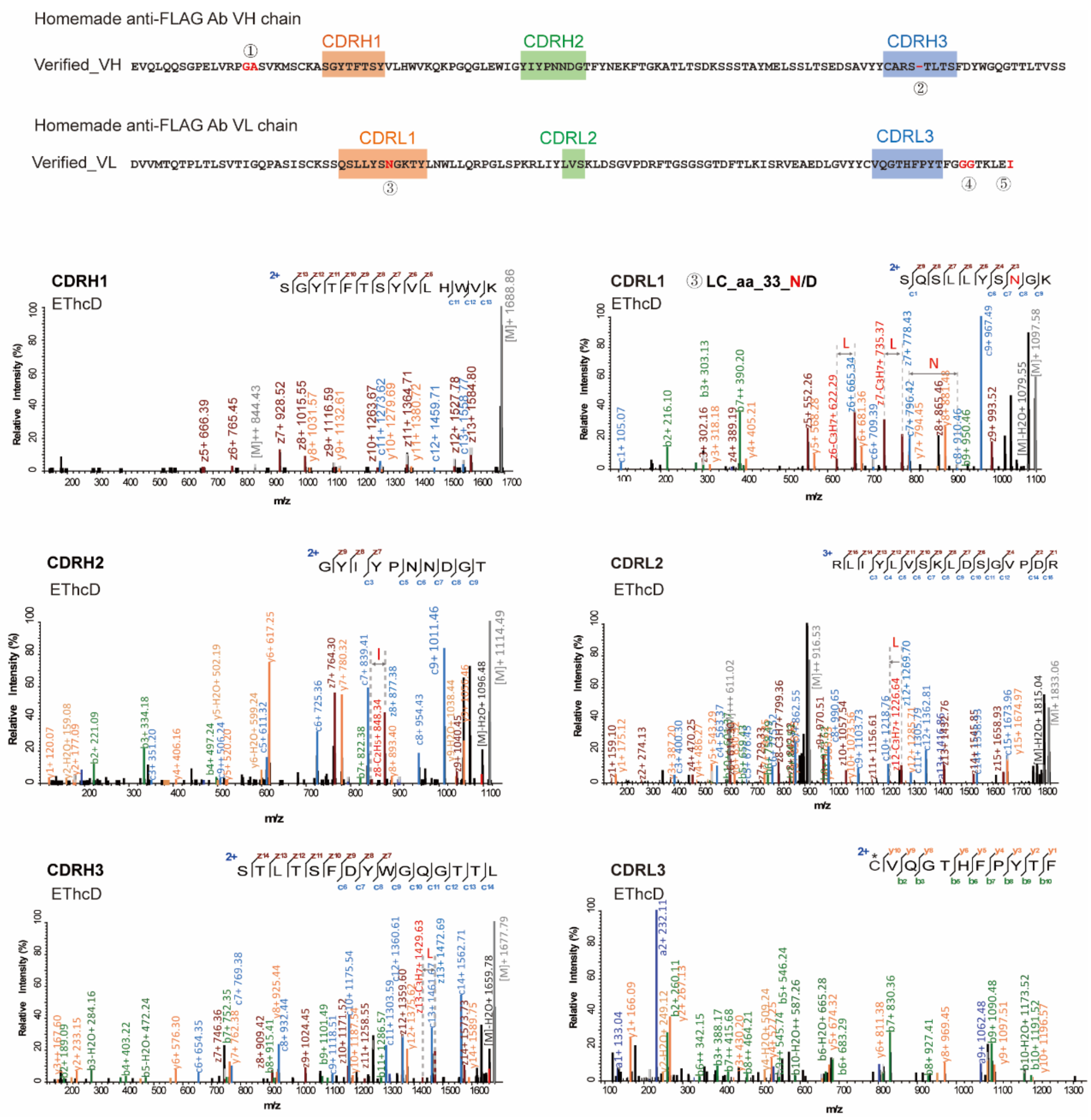


**Supplementary Figure 7.** Spectra supporting the reconstructed CDR sequences of the anti-HA antibody (TANA2). Carbamidomethylated C is labeled with * on the top in the peptide sequence displayed in the annotated MS2 spectra.

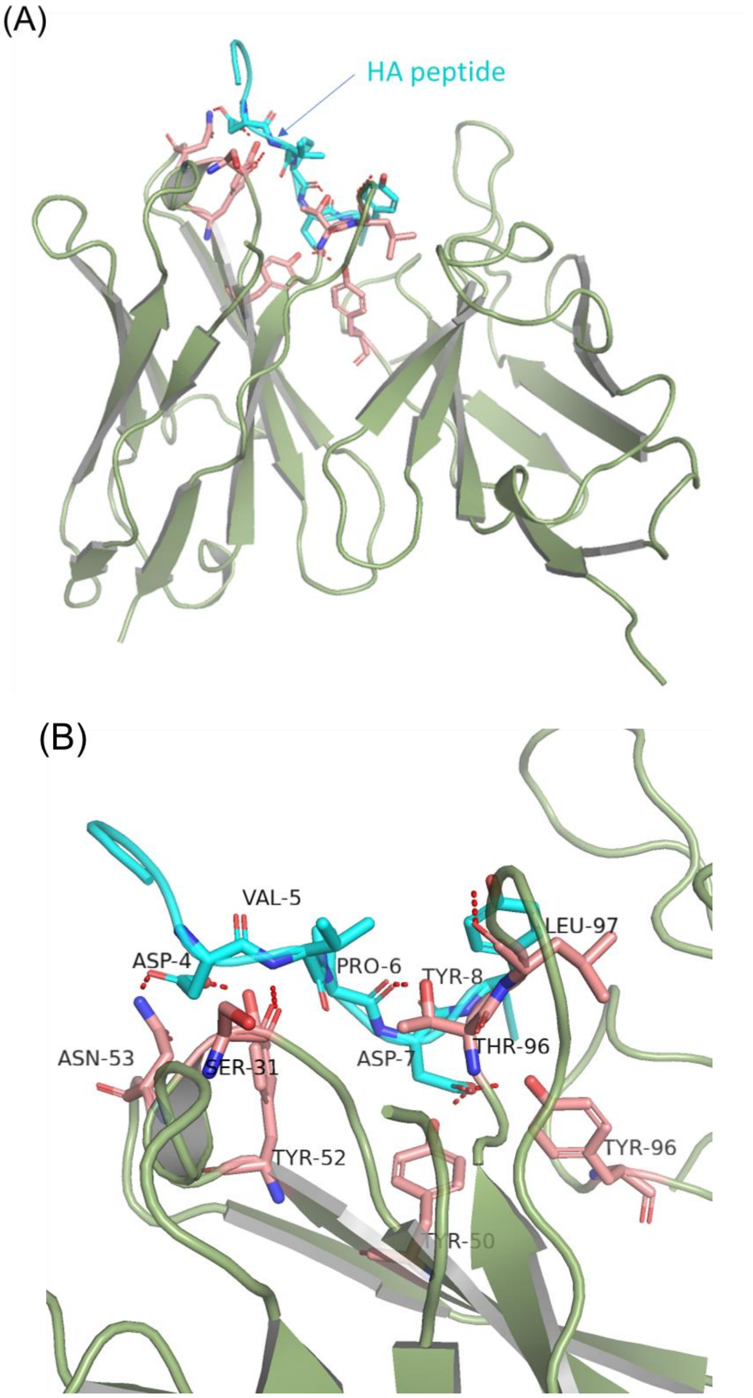


**Supplementary Figure 8.** A structure model of the anti-HA Ab in complex with the HA peptide, computed using RosettaAntibody 3. (A) The molecular docking results of TANA2 (shown in green) with the HA peptide (depicted in blue). (B) Amino acid residues lining the binding pocket are colored pink, and the key residues involved in hydrogen are marked.
